# Comprehensive profiling of genomic invertons in defined gut microbial community reveals associations with intestinal colonization and surface adhesion

**DOI:** 10.1101/2024.06.01.596983

**Authors:** Xiaofan Jin, Alice G. Cheng, Rachael Chanin, Feiqiao B. Yu, Alejandra Dimas, Marissa Jasper, Allison Weakley, Jia Yan, Ami S. Bhatt, Katherine S. Pollard

## Abstract

Bacteria use invertible genetic elements known as invertons to generate heterogeneity amongst a population and adapt to new and changing environments. In human gut bacteria, invertons are often found near genes associated with cell surface modifications, suggesting key roles in modulating dynamic processes such as surface adhesion and intestinal colonization. However, comprehensive testing of this hypothesis across complex bacterial communities like the human gut microbiome remains challenging. Metagenomic sequencing holds promising for detecting inversions without isolation and culturing, but ambiguity in read alignment limits the accuracy of the resulting inverton predictions. Here, we developed a customized bioinformatic workflow – PhaseFinderDC – to identify and track invertons in metagenomic data. Applying this method to a defined yet complex gut community (hCom2) across different growth environments over time using both *in vitro* and *in vivo* metagenomic samples, we detected invertons in most hCom2 strains. These include invertons whose orientation probabilities change over time and are statistically associated with environmental conditions. We used motif enrichment to identify putative inverton promoters and predict genes regulated by inverton flipping during intestinal colonization and surface adhesion. Analysis of inverton-proximal genes also revealed candidate invertases that may regulate flipping of specific invertons. Collectively, these findings suggest that surface adhesion and intestinal colonization in complex gut communities directly modulate inverton dynamics, offering new insights into the genetic mechanisms underlying these processes.

## Introduction

Bacteria in the human gut microbiome exist in complex communities with fluctuating dynamics [1–4] and spatial structure [5–7]. To successfully adapt to these environments, gut-associated strains harbor genomic inversions known as invertons that are capable of rapidly changing orientations and enable phase variation, i.e., phenotypic diversity across individual cells of the same strain [8–15]. At these invertons, invertase proteins bind specifically to inverted repeat (IR) regions in genomic DNA and mediate flipping of the intervening DNA sequence [16]. Previous work has analyzed closely related genomes and next generation sequencing datasets of microbial cultures to identify a large number of invertons in gut-associated microbes [14, 17], while computational approaches have yielded additional inverton predictions based on comparative genomics [18] and deep learning [19]. These invertons affect microbial phenotypes in multiple ways, for instance intergenic invertons could potentially regulate gene expression via inverton-embedded promoters, while gene-intersecting invertons can lead to new protein isoforms [17]. Invertons often regulate production of cell surface products such as exopolysaccharides, outer membrane proteins, and fimbriae [8–11, 13, 14], which are known to be associated with processes of gut colonization and surface adhesion [20–25]. However, the extent to which invertons modulate colonization and adhesion across a complex gut community remains unclear, as metagenomic approaches for comprehensive community-wide inverton profiling are limited by the general problem of sequence alignment ambiguity in communities with closely related strains [26].

We addressed these challenges by developing a bioinformatic workflow –PhaseFinderDC – to comprehensively profile invertons in defined communities of bacteria, based on the original PhaseFinder algorithm[14]. We then implemented this workflow on metagenomic samples of hCom2 – a defined yet complex community of bacterial strains modeled after the human gut[27] – grown across multiple conditions. These included: (i) isolate cultures of individual hCom2 strains [27, 28], (ii) fecal samples of gnotobiotic mice inoculated with hCom2 recovered from mice colonized over a total of 5 generations, and (iii) mixed cultures of hCom2 in various spatially structured in vitro environments [28] over a total of 6 passages. Our workflow successfully identified invertons in a majority of hCom2 strains and all eight represented phyla. For each identified inverton in each sample, we compared the proportion of sequencing reads supporting forward vs. reverse inverton orientations. Using this approach, we identified a subset of “directionally biased” invertons whose forward vs. reverse orientation probabilities are significantly different between growth conditions (e.g., isolate culture vs. *in vivo* mouse) and across timepoints (e.g., mouse generations or *in vitro* passages).

Categorizing identified invertons into inverton groups based on homology of their respective IR sequences, we applied motif enrichment analysis to identify motifs of IR and promoter sequences found in specific inverton groups. We identified gene families enriched in consistent orientations near invertons, highlighting cases where inverton-embedded promoters could potentially drive expression of “regulatable” downstream genes. We then used orientation of directionally biased invertons with regulatable genes to predict how surface adhesion and intestinal colonization dynamics are linked to expression of key bacterial genes, including surface-modifying genes such as those related to exopolysaccharide (EPS) biosynthesis in *Bacteroides*. Finally, we also highlight cases where specific invertase genes are enriched near specific inverton groups, potentially representing candidate invertases responsible for controlling inverton flipping. Together, these bioinformatic analyses provide a comprehensive community-wide characterization of inverton dynamics in a complex gut community, and point to key biological functions that modulate – and are modulated by – inverton-mediated phase variation.

## Results

### PhaseFinderDC detects invertons in defined communities with high sensitivity and specificity

Using the original PhaseFinder algorithm [14] as a starting point, we developed PhaseFind-erDC as a workflow to identify invertons in defined microbial communities for which reference genomes are available, with the ability to specifically discern invertons when closely related strains exist within the community. PhaseFinderDC was designed to take as input a concatenated reference genome database consisting of all strain genomes, and generate an alignment index by scanning this database for inverted repeats and compiling both forward and reverse orientation sequences for all inverted repeats (i.e., potential invertons), as per the original PhaseFinder algorithm. However as a modification, PhaseFinderDC also included sequences corresponding to genomic regions between inverted repeats so that the final sequence index covered the entire concatenated genome database and not merely the inverted repeat regions. The workflow then used bowtie2 to align short-read sequencing data to this index and quantify for each potential inverton the number of reads that align to the forward and reverse orientation references (Methods–Inverton detection using defined community sequencing data). Crucially, this comprehensive reference database allowed filtering of alignments by mapping quality (MAPQ) score and elimination ambiguous reads, thus addressing cases where highly related strains were present in the defined microbial community.

We implemented PhaseFinderDC on metagenomic samples derived from the hCom2 defined microbial community [27], which included several instances of closely related strains (Methods–Inverton detection using defined community sequencing data). We used a customized genome database generated by concatenating all hCom2 genomes (Supplementary Table S1, Fig. 1A). To benchmark the specificity of our updated PhaseFinder workflow, we used short-read sequencing data obtained from isolate cultures of each hCom2 strain (Supplementary Table S2) and quantified the number of reads that align to the forward and reverse orientation references for each potential inverton in hCom2. We called invertons based on reads counts from these isolate cultures if support for forward and reverse orientations had at least 5 reads each (Methods–Inverton detection using defined community sequencing data). We quantified the proportion of invertons called in the correct vs. incorrect strains, and found that our updated workflow produced 557 total calls, out of which 1 was mis-mapped (i.e., inverton called in a strain that was not the strain the sequencing data came from). This represented an improved specificity and sensitivity of inverton calls compared to the original PhaseFinder workflow which called 69 mismapped invertons out of 439 total calls with the same input read libraries and reference metagenome database (Fig. 1B, Supplementary Table S3). This improvement was especially dramatic amongst phylum Bacteroidota strains, wherein hCom2 has several instances of closely related strains. Indeed, closer examinination of this benchmarking that examined invertons called in Bacteroidota confirmed that mismapped invertons tended to occur between closely related strains (Fig. 1C,D), consistent with the known pitfall of read-stealing between closely related strains in metagenomic read alignment [26]. Altogether, these findings validated our updated workflow as a sensitive and specific approach for detecting invertons in defined bacterial communities.

**Fig. 1:**
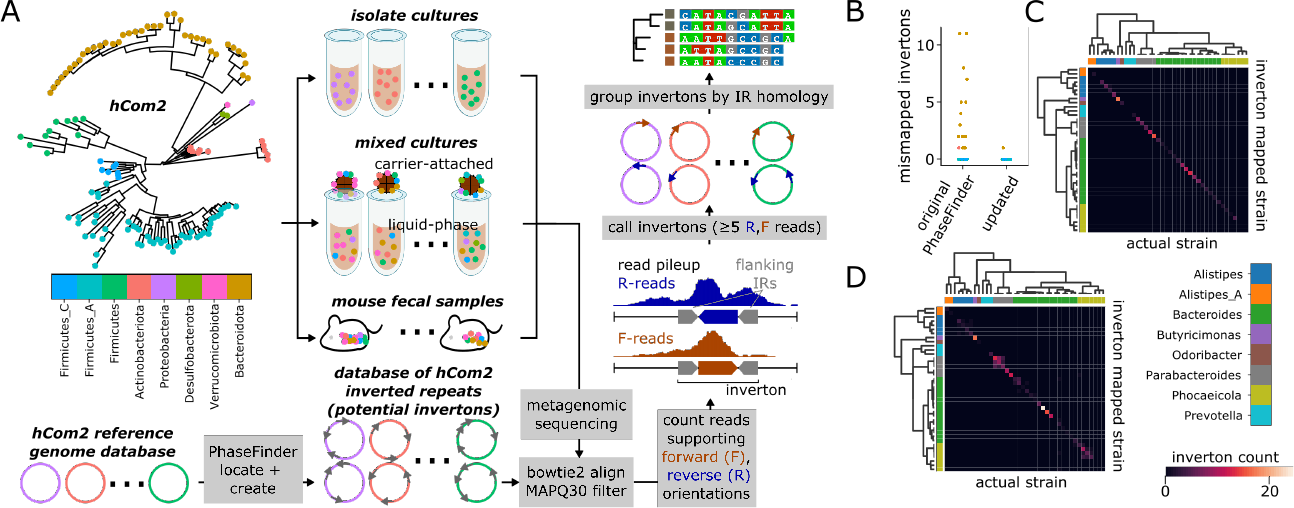
Customized workflow reliably detects invertons in metagenomes from defined communities (A) Flowchart of customized workflow to detect and group invertons in hCom2 from metagenomic sequencing data. (B) Benchmarking of workflow specificity based on pure isolate cultures - mismapped invertons refers to invertons called in one strain when using another strain’s sequencing library as input. (C) Heatmap of inverton counts among Bacteroides strains generated by updated PhaseFinder workflow , organized by actual strain (known based on isolate culture identity) and called inverton strain – off diagonal elements thus represent mismapped invertons. Strains are organized by phylogeny, margins correspond to genus. (D) Heatmap of inverton counts among Bacteroides strains generated by original PhaseFinder workflow, noting increase of off-diagonal (mismapped) calls between pairs of closely related strains.

Combining mapped reads from (i) isolate cultures of individual hCom2 strains, (ii) fecal samples of gnotobiotic mice inoculated with hCom2, and (iii) mixed in vitro cultures of hCom2 (Supplementary Table S2 for sequencing metadata), we applied our updated workflow – pooling forward and reverse orientation read counts across samples (Supplementary Table S4) – and detected 1837 invertons (Methods–Inverton detection using defined community sequencing data, Supplementary Table S5) across 99/125 strains in all 8 phyla present in the hCom2 community (Fig. 2A, Supplementary Table S5). Inverton counts per genome exhibited a wide range from *Mitsuokella multacida* DSM-20544 with 181 detected invertons, to 26 genomes without any detected invertons. For the 5 phyla in hCom2 with more than 2 representative strains, we found higher occurances of invertons in Bacteroidota with median 8.5 invertons per genome, Actinobacteriota with median 14 invertons per genome, and Firmicutes_C (primarily Negativicutes-like) with median 11 invertons per genome. Lower inverton counts were observed in Firmicutes_A (primarily Clostridia-like) with median 3 invertons per genome), and Firmicutes (primarily Bacillus-like) with median 0 invertons per genome. Comparing the genomic loci of identified invertons against those of predicted gene coding sequences (CDS), we found that just over half the invertons – 952/1837 – intersect a CDS, while the rest were intergenic (Supplementary Table S5). Invertons in Bacteroidota are primarily intergenic, while the opposite is true for Firmicutes_C. Invertons in Actinobacteriota, Firmicutes_A and Verrumicrobiota (with single representative strain *Akkermansia muciniphila*) ATCC-BAA-835 are roughly evenly split between intergenic and gene-intersecting (Supplementary Section S1).

**Fig. 2:**
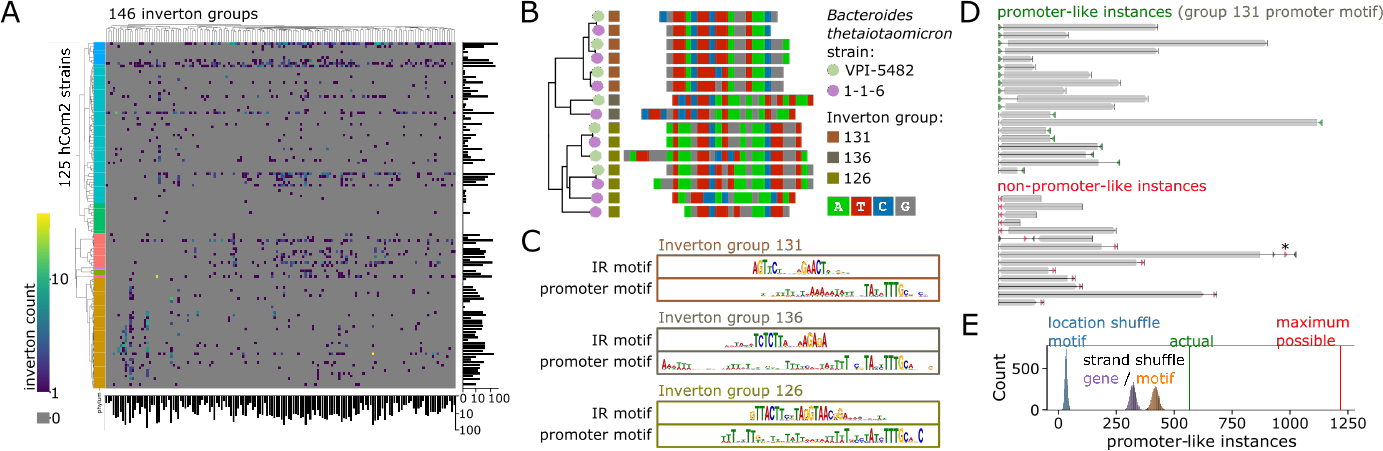
Enriched motifs detected in specific inverton groups grouped by IR sequence homology (A) Log-heatmap of inverton counts by hCom2 strain and by inverton group. hCom2 strains organized by phylogeny, phylum colors as in Fig. 1A. (B) Example of tree-building / grouping of identified invertons, subset on a group of 15 invertons in 2 *Bacteroides thetaiotaomicron* strains that were classified into groups 126, 131 and 136. (C) Promoter and IR motifs detected as enriched in inverton groups 126, 131 and 136. Promoter motifs found in each of these groups highly resembled previously described Bacteroides promoter motif [14, 29]. (D) Instances of promoter-like motif from inverton group 131 (motif 131-2) found in *Bacteroides thetaiotaomicron* VPI-5482 genome tended to be oriented upstream of nearest gene (grey). Instances are colored in green if consistent with promoter-like orientation, red otherwise. Asterisk marks instance of inverton embedded promoter that if flipped would be consistent with promoter-like orientation. (F) Across all of hCom2, 566 / 1217 total instances of motif 131-2 are consistent with promoter-like orientation – more than would be expected by random chance, based on random shuffling of motif loci, permuting motif strand, or permuting gene strand (10000 random samples tested for each case).

### Sequence homology in invertible regions enables categorization and motif enrichment analysis of detected invertons

We categorized the 1837 identified invertons into separate inverton groups (Fig. 2A) using homology of their respective IR sequences that flank each inverton. We performed a multiple sequence alignment (MSA) of all 1837 IR sequences, and used the resulting tree of sequences to cluster invertons into groups based on branch length thresholding (Fig. 2B, Supplementary Section S1, Methods–Inverton categorization using IR homology). The optimal branch length threshold was determined using IR sequence motif discoverability as a target metric (Supplementary Section S2, Methods–Inverton categorization using IR homology). This resulted in a total of 146 inverton groups (Figure 2A), with IR motifs identified for 114/146 inverton groups (Supplementary Table S6). Inverton groups ranged in size from 2 to 72 invertons with a median of 9 invertons per group. 29 out of the 146 groups had at least 10 invertons from a single phylum, with the largest group (inverton group 131) containing 68/72 invertons from Bacteroidota. Using max inverton count from a single phylum as a metric, we found that the 5 top inverton groups were all dominated by phylum Bacteroidota: after group 131, groups 126, 7, 136 and 143 contained 45/46, 42/46, 37/38, and 35/35 invertons respectively from Bacteroidota. We found that the IR motifs from these five groups matched closely to 5 previously published IR motifs found in Bacteroidota [14] (Supplementary Section S3). Furthermore, the IR motif for group 138 (which included 32/39 invertons from phylum Verrucomicrobiota, all from *Akkermansia muciniphila* ATCC-BAA-835) closely resembled the previously published IR motif in *Akkermansia muciniphila* [14]. This independent re-discovery of all 6 previously published IR motifs from Jiang *et al.* validated our inverton detection and grouping approach.

We also report a number of inverton groups with previously unpublished motifs across different phyla. For instance, focusing on the 29 inverton groups with at least 10 invertons from a single phylum, we found novel IR motifs in inverton groups 128, 137, 122, 48, and 68, which all comprised a majority of their invertons from phylum Bacteroidota, or inverton groups 139 and 125 which both comprised a majority of invertons from phylum Firmicutes_A (Supplementary Section S3). While many inverton groups appeared dominated by a single phylum, this was not always the case - as a counter-example, we also observed inverton group 145, which consisted of 20 invertons, 9 of which originated from phylum Bacteroidota (across 5 strains) and 10 of which originated from phylum Firmicutes_A (across 7 distinct strains). MSA of the 20 IR sequences in this group revealed a high degree of conservation even across phyla, and moreover leaves on the IR sequence MSA tree did not neatly cluster by phylum (i.e., IR sequences originating in Bacteroidota were not necessarily more similar to each other than those originating in Firmicutes_A, Supplementary Section S4). These findings suggest that group 145 invertons may potentially have spread across phyla via horizontal gene transfer.

Beyond IR motifs, we next applied motif enrichment search using MEME on full inverton sequences (as opposed to only their IR sequences) to identify motifs enriched in each inverton group (Methods–Motif detection and promoter prediction in invertons). We discovered a total of 266 motifs, spread across 126 inverton groups (Supplementary Table S7). Several identified motifs highly resemble previously reported inverton-associated promoters, consistent with the role of invertons turning gene expression on / off by changing promoter orientation[14]. For instance, we found motifs highly similar to a previously described Bacteroidota promoter motif [14, 29] independently enriched in 3 distinct Bacteroidota-dominated inverton groups (motifs 126-2, 131-2, and 136-2, Fig. 2C, Supplementary Section S3). Furthermore, we found that invertons in these three groups were differentially distributed between different Bacteroides strains (Supplementary Table S5), including closely related strains. As an example, we found that two strains of *Bacteroides thetaiotaomicron* present in hCom2 – VPI-5482 and 1-1-6, ANI estimate 98.8% using fastANI [30] – harbor 15 distinct invertons from groups 126, 131, and 136 (Fig. 2B).

An instance of a motif similar to the described Bacteroidota promoter [14, 29] was found in 14/15 of these invertons (i.e., either motif 126-2, 131-2, or 136-2). Using motif 131-2 as an example, we confirmed that across all of hCom2, instances of this motif tended to be found upstream of and on the same strand as their nearest gene (Fig. 2D), consistent with expectations for a promoter. We observed 566 out of 1217 total motif instances to exhibit this consistent upstream orientation, more than compared to random chance based on 10000 random samples each of (i) shuffling motif loci – median 32/1217 with consistent upstream orientation, (ii) permuting motif strand– median 417/1217 with consistent upstream orientation, and (iii) permuting gene strand – median 321/1217 with consistent upstream orientation (Fig. 2E). Meanwhile, we also discovered a motif enriched in sequences from inverton group 138 (which consists primarily – 32 out of 39 – of invertons from *Akkermansia*) that highly resembled the previously described *Akkermansia* promoter motif[14], which also exhibited promoter-like enrichment , though with a lower degree of confidence (Supplementary Section S5).

Beyond testing previously described promoter motifs, we also used random sampling bootstrap to detect new putative promoters. We scanned enriched motifs across all hCom2 metagenomes (Supplementary Table S8), and identified motifs whose detected instances were significantly enriched (p*<*0.001) on the same strand as and upstream of their nearest gene, meaning observed instances *>* 9990/10000 random samples for all three random sampling tests (shuffling motif loci, permuting motif strand, and permuting gene strand). This approach generated a catalog of 8 inverton-associated putative promoter motifs, which included both previously described promoter motifs as well as several previously undescribed motifs (Supplementary Section S5). Note that based on the (p*<*0.001) cutoff used, the previously described Bacteroidota promoter motif was counted as a putative promoter motif (listed three times independently as motif 126-2, 131-2 and 136-2), but not the previously described *Akkermansia* motif (motif 138-2) [14] (Supplementary Section S5). In addition to putative promoters, we also found motifs enriched on the same strand as and downstream of their nearby gene (Supplementary Section S5), which may be indicative of sequence features associated with transciptional termination or post-transcriptional modification. Collectively, these findings demonstrated the utility of our community-wide bioinformatic search as an approach for motif and promoter discovery.

### Genomic proximity links distinct inverton groups to specific gene families

We next sought to identify gene families that are enriched in regions proximal to (within +/-5kb) or directly intersecting identified invertons. As promoters are known to often be embedded within invertons[14], we further split non-inverton-intersecting genes between ‘regulatable’ genes which could be driven by a promoter in their associated inverton (i.e., 5’-end is proximal to inverton for the given gene and all genes between given gene and inverton), and ‘non-regulatable’ genes that are nevertheless proximal to invertons (Supplementary Table S9).

We uncovered numerous cases of inverton-group specific gene family enrichment for both ‘regulatable’ and ‘non-regulatable’ genes, as well as genes that directly intersect invertons (Supplementary Table S10). For instance, UpxY family transcription antiterminator was significantly enriched as regulatable genes (Fig. 3A,B) near invertons in Bacteroidota-dominated groups 126, 144, 136 and 137, suggesting possible roles in regulation of exopolysaccharide biosynthesis [9]. Other enriched regulatable genes included many previously described hits related to cell surface products such as TonB-dependent receptor and RagB/SusD family nutrient uptake outer membrane protein (inverton group 131), fimbrillin family protein (inverton groups 137, 112, 122), and PEP-CTERM sorting domain-containing protein (inverton group 138).

**Fig. 3:**
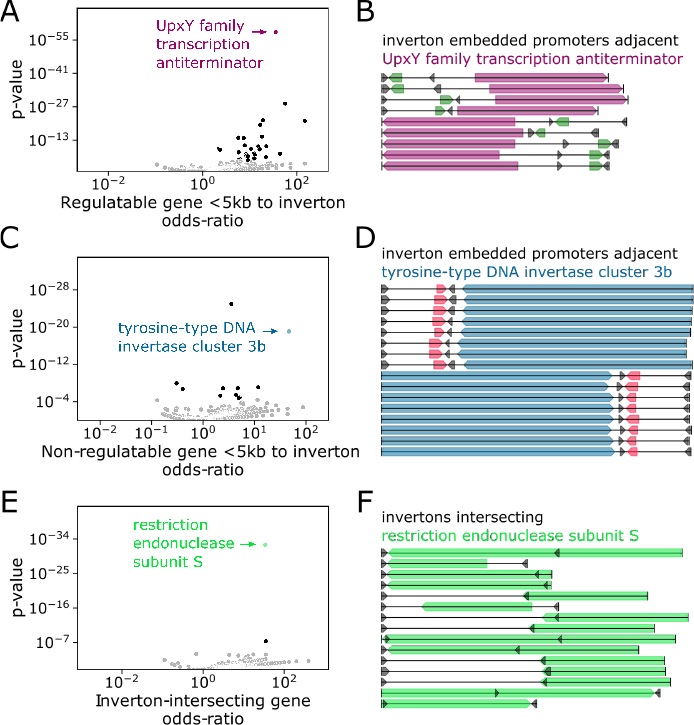
Cell surface products and invertase genes are enriched with consistent orientation near specific inverton groups (A) Volcano plot of regulatable gene families enriched near invertons based on odds ratio of being observed within 5kb of an inverton, relative to rest of the hCom2 genomes. Grey points are non-significant based on Bonferroni p-value correction. (B) Genome diagram of all instances of UpxY family transcription antiterminator genes near invertons containing previously described Bacteroides promoter motif [14, 29] – coding sequences annotated with this gene function (marked in purple) are consistently oriented in a way that can be regulated by promoter motif (marked in green) upon inverton flipping. Flanking inverton IR regions marked in black. (C) Volcano plot of non-regulatable gene families enriched near invertons based on odds ratio of being observed within 5kb of an inverton, relative to rest of the hCom2 genomes. Grey points are non-significant based on Bonferroni p-value correction. (D) Genome diagram of all instances of tyrosine-type DNA invertase cluster 3b genes near invertons containing previously described Bacteroides promoter motif [14, 29] – coding sequences annotated with this gene function (marked in blue) are consistently oriented in a way that cannot be regulated by promoter motif (marked in red) upon inverton flipping. Flanking inverton IR regions marked in black. (E) Volcano plot of gene families enriched for direct inverton intersection, based on odds ratio of being directly overlapping an inverton, relative to rest of the hCom2 genomes. Grey points are non-significant based on Bonferroni p-value correction. (F) Genome diagram of all instances of restriction endonuclease subunit S genes near invertons from group number 145. Coding sequences annotated with this gene function (in green) consistently overlap inverton loci. Flanking inverton IR regions marked in black.

Meanwhile, enriched non-regulatable genes included several invertase families such as tyrosine-type DNA invertase cluster 3b (Fig 3C,D), which was enriched in inverton group 126, as well as tyrosine-type recombinase/integrase (inverton groups 139, 145) and sitespecific integrase (inverton groups 136, 126, 137, 48, 116). The presence of cell surface products and invertase genes near invertons aligned with previous reports of similar gene enrichment in gut microbes[14]. Our own results further indicated that invertases near invertons – which are often considered likely candidates for controlling inverton flipping [16] – are generally non-regulatable and thus unlikely to themselves be regulated by invertonembedded promoters. A potential exception however was found in inverton group 145, with tyrosine-type recombinase/integrase gene family found to be enriched in both regulatable as well as non-regulatable orientations near group 145 invertons.

Finally we also observed instances of gene families enriched for direct intersection with inverton sequences. For instance, invertons from group 145 were significantly enriched with members of the restriction endonuclease subunit S gene family (Fig. 3E,F). This observation aligns with previous reports of restriction enzymes with switchable specificity where recombination and inversion at enzyme coding sequences leads to production of different isoforms of enzyme protein with different specificities [31–34]. In addition to enrichment of restriction endonuclease subunit S and tyrosine-type recombinase/integrase gene families, inverton group 145 was also enriched for the relaxase/mobilization nuclease domain-containing protein and plasmid mobilization relaxosome protein MobC gene families genes, lending further support to the the idea that this group of invertons may have spread via HGT.

### Genomic proximity links distinct inverton groups to specific invertases

We next explored whether a bioinformatic approach could link identified inverton groups to specific groups of invertase genes, based on the idea that such genes are often located near the invertons they regulate [16]. Gene annotation detected 3932 invertase genes amongst hCom2 genomes (Supplementary Table S11). We used multiple sequence alignment and clustering to group these invertase genes into 176 invertase groups based on sequence homology (Fig. 4A, Methods – Genome annotation, invertase detection and categorization, Supplementary Section S6). For each of the 126 identified inverton groups, we systematically checked for each of the 176 invertase groups whether the corresponding invertases are enriched in the proximity of the corresponding invertons.

**Fig. 4:**
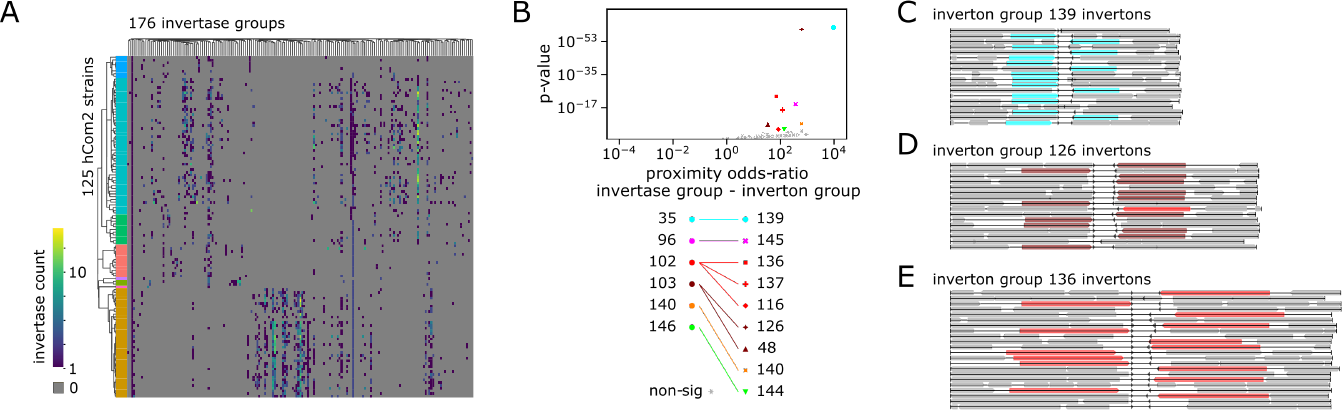
Specific invertase groups are enriched near specific inverton groups (A) Log-heatmap of invertase counts by hCom2 strain and by invertase group. hCom2 strains organized by phylogeny, phylum colors as in Fig. 1A. (B) Volcano plot of inverton-group to invertase-group pairs, based on odds ratio of invertase in given invertase group being observed within 5kb of an inverton in given inverton group. Grey points are non-significant based on Bonferroni p-value correction. Inverton and invertase groups are specified by marker color and shape respectively, with significant links specified in the legend. (C) Genome diagram of regions surrounding group 139 invertons, with invertase group 35 genes highlighted in blue – for simplification if multiple invertons of this group are found in the same genome, a single representative is plotted. Nearly all invertons in this group have a nearby group 35 invertase. (D) Genome diagram of regions surrounding group 126 invertons, with invertase group 103 genes highlighted in dark red – for simplification if multiple invertons of this group are found in the same genome, a single representative is plotted. (E) Genome diagram of regions surrounding group 136 invertons, with invertase group 102 genes highlighted in light red – for simplification if multiple invertons of this group are found in the same genome, a single representative is plotted.

Counting an invertase gene as proximal if it intersects, or is within 5kb of the inverton, we identified inverton group - invertase group pairs with significantly (Fisher’s Exact p*<*0.05 with Bonferroni correction) enriched proximal invertase counts (Supplementary Table S12). This approach revealed 9 such pairs (Fig. 4B, Methods), representing a total of 135 inverton-invertase examples from 52 different strains including members of both Clostridia and Bacteroidia classes (Supplementary Table S11). The most significant link between such pairs was between inverton group 139 – a group of 33 invertons (32 of which are from class Clostridia) – and invertase group 35, a group of genes belonging to the tyrosine-type recombinase/integrase family found in classes Clostridia and Bacteroidia (Fig 4B,C). Meanwhile, among the Bacteroidota-dominated inverton groups, group 136, 137, and 116 had significant links to invertase group 102, while inverton groups 126 and 48 were instead linked to invertase group 103 (Fig 4B,D,E). By contrast, other large Bacteroides-dominated inverton groups such as 131 were not significantly linked to any invertase groups (Fig. 4B, Supplementary Section S7). These associations between inverton and invertase groups indicates potential co-evolution of invertase and IR sequences, with instances of multiple inverton groups linked to a single invertase group potentially further suggesting some degree of flexibility in the ability of invertase proteins to control flipping across inverton recognition IR sites.

### Differential analysis between metagenomic sample types indicates inverton-modulated gene expression changes associated with surface adhesion and gut colonization

To investigate how processes of bacterial surface adhesion and gut colonization are linked to inverton orientations at a community-wide level, we compared inverton orientation probabilities between different sample types (Supplementary Table S4). We first identi-fied associations with gut colonization by searching for invertons where our bioinformatic workflow yields a different proportion of sequencing reads supporting the forward and reverse orientations between (i) samples from isolate cultures of individual hCom2 strains and (ii) fecal samples of gnotobiotic mice inoculated with hCom2 (Fig 5A,B), recovered from mice colonized over a total of 5 generations. As an example, we found an inverton in inverton group 126 – B-th-VPI-5482 0:4315126-4315146-4315394-4315414 – which had read support for both forward and reverse orientations in *Bacteroides thetaiotaomicron* VPI-5482 isolate culture, but only support for the reverse orientation in mouse-stool sequencing samples (Fig. 5A), a trend that was consistent across 69 mouse stool samples and 3 isolate cultures (Fig. 5B). All told we identified 224 directionally biased invertons (across 53 strains) whose forward and reverse read proportions were significantly (Fisher’s Exact p*<*0.05 with Bonferroni correction) different between pooled isolate culture and pooled mouse gut fecal samples (Fig. 5A, Methods, Supplementary Table S13). Applying the same approach we next identified associations specifically with surface adhesion by comparing (iii-a) mixed in vitro cultures of hCom2 as a surface attached community using mucin-agar carriers as a synthetic surface, and (iii-b) corresponding mucin-agar carrier cultures of hCom2, instead sampled from the liquid-phase. This yielded 38 directionally biased invertons (across 16 strains) whose orientation probabilities were significantly different between mucin-agar surface-attached and liquid-phase samples (Supplementary Table S13).

**Fig. 5:**
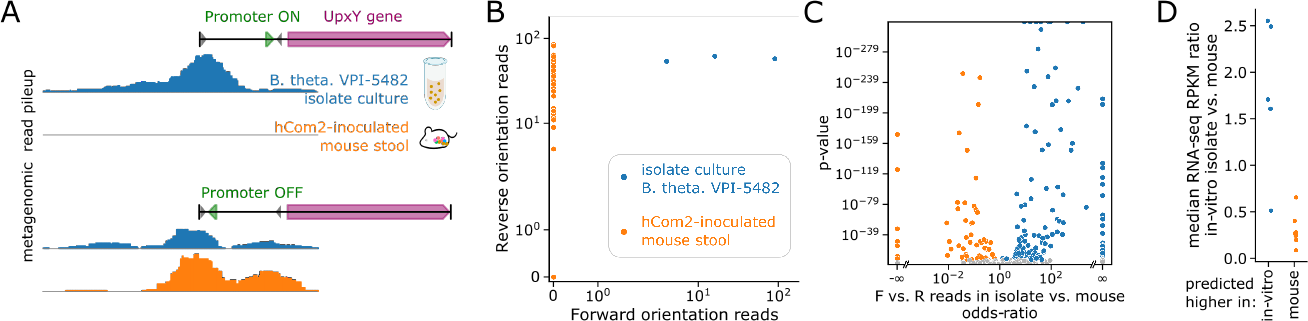
Inverton orientation is modulated by gut colonization and drives differential gene expression (A) Example of inverton in *Bacteroides thetaiotaomicron* VPI-5482 from inverton group 126 (B-th-VPI-5482 0:4315126-4315146-4315394-4315414) with different forward and reverse read counts depending on growth condition. While roughly equal forward and reverse read support is observed when cultured in vitro, mouse stool derived samples only show reverse orientation read support. (B) Forward vs. reverse read count scatterplot of inverton in (A), demonstrating that reverse orientation enrichment is observed consistently across all mouse samples. (C) Volcano plot of invertons with directional preference, based on odds ratio of forward vs. reverse reads count for in vitro isolate vs. mouse samples. (D) Median RPKM in-vitro vs. mouse ratios for 12 *Bacteroides thetaiotaomicron* VPI-5482 genes predicted to be differentially expressed between in vitro isolate vs. mouse samples based on inverton orientation. Values greater than 1 correspond to in vitro upregulation relative to mouse. Blue and orange dots represent genes predicted to be upregulated in vitro and in mouse respectively.

Cross-referencing these directionally biased invertons against (1) our catalog invertonembedded putative promoters (Supplementary Section S5, Supplementary Table S8) and (2) locations and orientations of regulatable gene CDSs adjacent to these invertons (Supplementary Table S9), we generated lists of genes whose expression we predicted to be modulated either up or down by inverton-flipping during either gut colonization (comparing isolate culture vs. mouse stool samples) and surface adhesion (comparing carrier-attached vs. supernatant mixed culture samples) (Methods Gene expression prediction, Supplementary Table S14). We found a number of significant (Fisher’s Exact p*<*0.05 with Bonferroni correction) gene families related to cell surface products (Supplementary Table S15). Some, such as the SLBB domain-containing protein and SH3 beta-barrel fold-containing protein families appeared to be consistently up-regulated in mouse stool compared to isolate culture (Supplementary Table S15), suggesting higher expression during gut colonization. Meanwhile, other gene families such as UpxY family transcription antiterminator and polysaccharide biosynthesis/export family protein – both of which are linked to EPS production in Bacteroidota [9] – appeared to be enriched for both down as well as upregulation (Supplementary Table S15) during gut colonization (i.e., more of these genes are up-regulated than would be expected by random chance, and also more of these genes are down-regulated than would be expected by random chance), consistent with the notion that cells may be actively remodeling their surface EPS content by turning off production of certain types in favor of others [9]. Comparing carrier-attached vs. supernatant cultures, we found the FimB/Mfa2 family fimbrial subunit gene family was consistently up-regulated on carrier attached cultures (Supplementary Table S15), consistent with the known role of these genes in adhesion and biofilm formation [35].

Next, we sought to use publicly available transcriptomic data to validate some of the differential gene expression predictions we made based on directional enrichment of promoter-embedded invertons. We leveraged a recently published RNA-seq dataset of *Bacteroides thetaiotaomicron* VPI-5482 [36], with read libraries derived from both in vitro culture as well as mouse samples (Table S16). Based on our analysis, we computationally predicted a total of 12 differentially expressed genes in *Bacteroides thetaiotaomicron* VPI-5482 when comparing between isolate in vitro culture and growth in mouse, 5 of which we predict to be upregulated in vitro and 7 of which we predict to be upregulated in mouse. To validate these predictions, we calculate the Median RNA-seq Reads Per Kilobase per Million mapped reads (RPKM) for these 12 genes in both mouse and in vitro RNA-seq data (Table S17). These calculations reveal that the median RPKM in-vitro vs. mouse ratios are significantly higher (p=0.0047, two-sided Mann-Whitney U test) in genes we predicted as in-vitro upregulated than in genes we predicted as mouse upregulated (Fig. 5D), providing preliminary validation for our approach. Collectively, these findings confirmed that surface adhesion and intestinal colonization in complex gut communities directly modulate inverton flipping, and predicted how the expression of key genes are modulated in this process.

### Longitudinal analysis across *in vivo* and *in vitro* timepoints

In addition to performing differential analysis between different metagenomic sample types (e.g., isolate culture vs. mouse stool), we also analyzed longitudinal samples across timepoints to better understand inverton dynamics both *in vivo* and *in vitro*. For *in vivo* analysis, we compared samples across mouse generations, and searched for invertons whose forward-vs.-reverse orientations exhibited significant (Fisher’s Exact p*<*0.05 with Bonferroni correction) differences between early and late generation mice (Methods – Longitudinal analysis of inverton orientation across timepoints). We additionally calculate at each timepoint the F.-vs.-R. inversion ratio – defined as reverse over total read counts (R/(R+F)).

Using this approach, we identified a total of 123 invertons (Table S18) across 34 strains that exhibited time-dependent behavior, such as an inverton in *Akkermansia muciniphila* ATCC-BAA-835 from inverton group 138 located near two autotransporter domain containing proteins (A-mu-ATCC-BAA-835 0:2092093-2092109-2092267-2092283). The F.-vs.-R. inversion ratio for this inverton trended from nearly universal reverse orientation in first generation (SC1) mice to majority forward orientation by fifth generation (SC5) mice (Fig., 6A, Supplementary Section S8). Curiously, we also found this type of trend in some – but not all – other group 138 invertons from *Akkermansia muciniphila* (Supplementary Section S9), suggesting the existence of additional layers of regulatory control in determining inverton flipping dynamics beyond simple IR sequence recognition.

Our analysis also revealed a wide range of timescales of inverton dynamics, sometimes even within a single bacterial strain. For instance, within *Bacteroides cellulosilyticus* DSM-14838, we observed two distinct invertons – B-ce-DSM-14838 0:4484223-4484241-4484427-4484445 from inverton group 143 and B-ce-DSM-14838 0:137243-137258-137893-137908 from inverton group 131 – that both started from nearly complete forward orientation in SC1 mice and trended toward increasing reverse orientation but with the former doing so at a markedly more rapid rate (Fig. 6B,C). The presence of nutrient uptake genes near the former and exopolysaccharide biosynthesis genes near the latter suggested these two invertons may be responsible for regulating different biological functions (Supplementary Section S8).

**Fig. 6:**
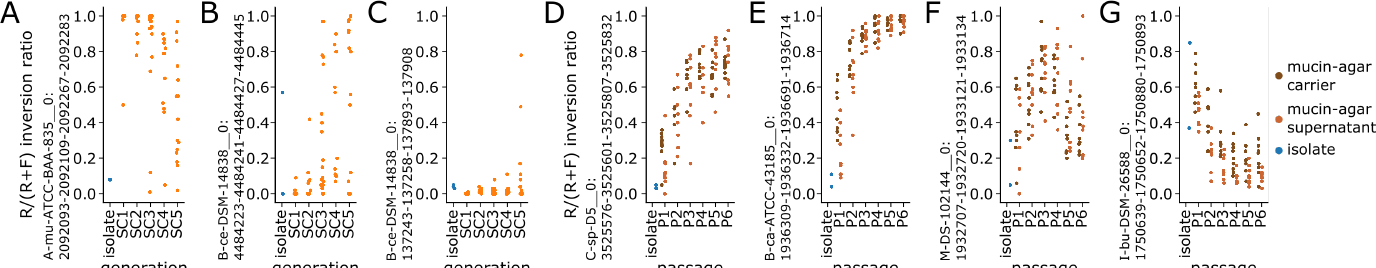
Invertons exhibit dynamic flipping across *in vivo* mouse generations and ***in vitro*** culture passages (A) Example of inverton in *Akkermansia muciniphila* ATCC-BAA-835 from inverton group 138 (A-mu-ATCC-BAA-835 0:2092093-2092109-2092267-2092283) whose inversion ratio (R/(R+F), so higher scores correspond to more reverse orientation reads) exhibits a dynamic trend across mouse generations, from nearly universal reverse orientation in first generation (SC1) mice, but shifts toward forward orientation in later generations. (B) Example of inverton in *Bacteroides cellulosilyticus* DSM-14838 from inverton group 143 (B-ce-DSM-14838 0:4484223-4484241-4484427-4484445) whose inversion ratio shifts from nearly universal forward orientation in first generation (SC1) mice toward reverse orientation in later generations. (C) Example of inverton in *Bacteroides cellulosilyticus* DSM-14838 from inverton group 131 (B-ce-DSM-14838 0:137243-137258-137893-137908) whose inversion ratio shifts from nearly universal forward orientation in first generation (SC1) mice toward reverse orientation in later generations, with a markedly slower trend than (B). (D) Example of inverton in *Clostridium sp.* D5 from inverton group 139 (C-sp-D5 0:3525576-3525601-3525807-3525832) whose inversion ratio shifts from mostly forward orientation in passage 1 cultures (P1) toward reverse orientation in later passages. (E) Example of inverton in *Bacteroides caccae* ATCC-43185 from inverton group 131 (B-ca-ATCC-43185 0:1936309-1936332-1936691-1936714) whose inversion ratio shifts rapidly from mostly forward orientation in passage 1 cultures (P1) toward nearly complete reverse orientation by P4/P5. (F) Example of inverton in *Megasphaera* DSMZ-102144 from inverton group 51 (M-DS-102144 0:1932707-1932720-1933121-1933134) whose inversion ratio shifts from mostly forward orientation in passage 1 cultures (P1) toward reverse orientation, before stabilizing at a roughly equal mix of reverse and forward orietation reads. (G) Example of inverton in *Intestinimonas butyriciproducens* DSM-26588 from inverton group 139 (I-bu-DSM-26588 0:1750639-1750652-1750880-1750893) whose inversion ratio shifts from mostly reverse orientation in passage 1 cultures (P1) toward forward orientation in later passages, with consistently higher inversion ratios in mucin-agar carrier cultures (dark brown) than corresponding supernatant cultures (light brown).

Applying this early-vs.-late enrichment approach with mixed *in vitro* cultures, we also identified invertons whose forward-vs.-reverse orientations were associated with changes across passage timepoints (Table S18), such as an inverton in *Clostridium sp.* D5 from inverton group 139 (C-sp-D5 0:3525576-3525601-3525807-3525832) which exhibits a dynamic shift from mostly forward orientation in passage 1 cultures (P1) toward reverse orientation in later passages (Fig. 6D). As with mouse generational data, we also observed a range of timescales, with an inverton in *Bacteroides caccae* ATCC-43185 from inverton group 131 (B-ca-ATCC-43185 0:1936309-1936332-1936691-1936714) exhibiting particularly a rapid shift from mostly forward orientation in passage 1 cultures (P1) toward nearly complete reverse orientation by P4/P5 (Fig. 6E). Note additionally that timescales involved in *in vitro* cultures are inherently much shorter than mouse generational data as culture passage intervals are 3 days apart, compared with months between mouse generations.

We also observed invertons with dynamics that do not appear to converge toward fully reverse nor fully forward orientation, such as one in *Megasphaera* DSMZ-102144 from inverton group 51 (M-DS-102144 0:1932707-1932720-1933121-1933134) which exhibits an early shift toward reverse orientation, before stabilizing at a roughly equal mix of reverse and forward orientation reads at later passages (Fig. 6F). Finally, we also observed invertons whose *in vitro* dynamics appeared to be dependent on surface adhesion, such as inverton in *Intestinimonas butyriciproducens* DSM-26588 from inverton group 139 (I-bu-DSM-26588 0:1750639-1750652-1750880-1750893) which exhibited increasing forward orientation overall, but with consistently higher reverse orientation in surface-attached mucin-agar carrier cultures than corresponding supernatant cultures (Fig. 6G). Carrier-vs.-supernatant dependent dynamics were also observed in a collection of 19 invertons (across 13 different inverton groups) all from the same strain of *Clostridium sp.* D5 that consistently exhibited modest levels of reverse orientation reads in carrier cultures, with virtually no reverse orientation reads in supernatant cultures (Supplementary Section S10). These findings that invertons from different inverton groups (i.e., with disparate IR sequences) can exhibit highly similar trends between surface-attached vs. supernatant cultures over time, while those with highly similar IR sequences can exhibit disparate trends across mouse generations (Supplementary Section S9) – reinforce the idea that invertons produce a wide range of dynamic behaviors during surface adhesion and gut colonization, and that these dynamics are likely mediated by additional layers of regulatory control beyond IR sequence recognition alone.

## Methods

### hCom2 mouse generational experiment and metagenomic sequencing

Three male-female pairs of germ free mice were colonized with hCom2 and bred for three generations. Male and female pups were weaned at age 4 weeks and housed in separate cages. For each generation three male-female pairs were co-housed and bred to generate the next generation. Extra pups that were not used for breeding were housed in separate cages by gender. Pups from the Parental generation were labeled as SC1, F1 generation labeled as SC2, F2 generation SC3, and F3 generation as SC4. At the beginning of each month, fecal sampling was performed and the stool was sequenced. Metagenomic sequencing was performed as previously described [27]: genomic DNA was extracted from pellets using the DNeasy PowerSoil HTP kit (Qiagen) and quantified in 384-well format using the Quant-iT PicoGreen dsDNA Assay Kit (Thermofisher). Sequencing libraries were generated in 384-well format using a custom low-volume protocol based on the Nextera XT process (Illumina). The concentration of DNA from each sample was normalized to 0.18 ng/µL using a Mantis liquid handler (Formulatrix). If the concentration was <0.18 ng/µL, the sample was not diluted further. Tagmentation, neutralization, and PCR steps of the Nextera XT process were performed on a Mosquito HTS liquid handler (TTP Labtech), leading to a final volume of 4 µL per library. During PCR amplification, custom 12-bp dual unique indices were introduced to eliminate barcode switching, a phenomenon that occurs on Illumina sequencing platforms with patterned flow cells. Libraries were pooled at the desired relative molar ratios and cleaned up using Ampure XP beads (Beckman) to achieve buffer removal and library size selection. The cleanup process was used to remove fragments < 300 bp or > 1.5 kbp. Final library pools were quality-checked for size distribution and concentration using a Fragment Analyzer (Agilent) and qPCR (BioRad).

Sequencing reads were generated using a NovaSeq S4 flow cell or a NextSeq High Output kit, in 2x150 bp configuration. 5-10 million paired-end reads were targeted for isolates and 20-30 million paired-end reads for communities.

#### Inverton detection using defined community sequencing data

We developed a modified Phasefinder [14] workflow – PhaseFinderDC – to detect invertons from metagenomic read libraries derived from mixtures of hCom2 strains. A custom hCom2 reference database was generated by concatenating microbial genome sequences for all 125 strains in hCom2. As in the original Phasefinder workflow, EMBOSS einverted was then used to locate inverted repeat sequences and thus generate a list of potential invertons. Based on this list, we then created an augmented genomic reference containing both forward and reverse orientation sequences for each potential inverton. This augmented reference then served as the database against which metagenomic sequencing reads are aligned. Here we made a modification to the original Phasefinder workflow for PhaseFinderDC to include all genomic sequences located between potential invertons – in addition to forward / reverse sequences of potential invertons – to aid in filtering of ambiguously aligned reads (discussed further below).

Metagenomic read libraries derived from mouse stool samples were pre-processed using Biobakery kneaddata [37] to remove mouse DNA reads – this was skipped for samples derived from pure single-strain and mixed community in vitro cultures. Metagenomic read alignment of read libraries from was then carried out using bowtie2 instead of bowtie (as per original Phasefinder), given that the majority of sequencing reads used in our analyses exceeded 100bp in length. Using mapping quality (MAPQ) scores reported by bowtie2, PhaseFinderDC filtered out ambiguously aligned reads by removing any alignments with scores below 30. For each potential inverton, counts of read alignments unambiguously supporting forward and reverse orientations based on paired-end orientation (Pe_F, Pe_R) and based on directly spanning inversion junction (Span_F, Span_R) were enumerated as per the original Phasefinder workflow for each sample. For each potential inverton, read counts were pooled across isolate culture samples from the given strain, as well as all hCom2 mouse and in vitro mixed culture samples. We called invertons if forward and reverse orientations are supported by at least 5 read alignments each after pooling, based on both paired-end orientation as well as direct span (i.e., required pooled *Pe*_*F >*= 5*, Pe*_*R >*= 5*, Span*_*F >*= 5*, Span*_*R >*= 5). This threshold is similar to that suggested by the original PhaseFinder publication [14] which used *Pe*_*F >*= 5*, Pe*_*R >*= 5*, Span*_*F >*= 3*, Span*_*R >*= 3. The original publication also used a R/(R+F)*>*0.01 cutoff, which we omitted here to enable capturing of rarely flipped invertons, compensating instead with a slightly more stringent *Span*_*F >*= 5*, Span*_*R >*= 5 cutoff. Downstream analysis such as calculation of inversion ratios for each called inverton for each sample used Pe_F, Pe_R counts.

### Inverton categorization using IR homology

IR sequences for all 1832 called invertons were used to generate a multiple sequence alignment using Clustal-omega [38]. Based on this MSA, we used TreeCluster [39] to cluster invertons into distinct groups based on IR sequence homology. We tested a range of tree-distance thresholds (T=0.01 to 0.99 in increments of 0.01) for generating separate groups, with smaller thresholds generating more (smaller, more closely related) groups of invertons. We then used MEME [40] on IR sequence groups to search for enriched sequence motifs applying -mod zoops -nmotifs 1000 -minw 6 -maxw 100 -objfun classic -revcomp -markov_order 0 -evt 0.05 parameters, counting the total number of groups with a detected motif for each tested tree-distance threshold. We then selected a tree-distance cutoff of T=0.60 as it maximized the number of groups for which a motif was detected, generating a total of 146 groups of invertons out of which 114 MEME was able to detect an enriched sequence motif in the IR sequences (Supplementary Section S2).

### Motif detection and promoter prediction in invertons

For each of the 146 identified groups of invertons, we used MEME to identify motifs enriched within inverton sequences applying -mod anr -nmotifs 1000 -minw 6 -maxw 100 -objfun classic -revcomp -markov_order 0 -evt 0.05 parameters, now using the full inverton sequences, as opposed to only the IR sequences. Significantly enriched motifs were then scanned using fimo across all hCom2 genomes, applying a 10*^−^*^8^ p-value cutoff to account for the large sequence database size (4.67*·*10^8^bp total). For each motif instance detected by fimo, we used bedtools closest [41] to identify location and orientation of the closest associated coding sequence. For each motif, we then counted the number of fimo-detected instances that were consistent with those of a promoter, that is to say upstream of its nearest gene, on the same strand. We compared this number against (i) 10000 random samples where the locations of motif instances were shuffled using bedtools shuffle [41], (ii) 10000 random samples where the strand orientations of motif instances were permuted, and (iii) 10000 random samples where the strand orientations of hCom2 genes were permuted. Motifs whose actual count of promoter-consistent instances exceeded 9990/10000 (>99.9%ile) of all three random samples were identified as putative promoter motifs. Note that as we did not know *a priori* whether the motif or its reverse complement represented a putative promoter element, we performed these tests for all identified motifs and their reverse complements, reporting the version oriented on the same strand as the nearest gene.

### Gene annotation and enrichment analysis near invertons

All hCom2 genomes were annotated using NCBI PGAP pipeline [42] version 2023-05-17.build6771. We identified enrichment of specific annotations near invertons by counting for each annotation the number of instances (i) gene with given annotation is located within +/-5kb window of any detected inverton, (ii) gene with different annotation is located within +/-5kb window of any detected inverton, (iii) gene with given annotation is not located within +/-5kb window of any detected inverton, and (iv) gene with different annotation is not located within +/-5kb window of any detected inverton. For each gene annotation, we compiled these four counts into a 2x2 contingency table, and identified annotations significantly (Fisher’s Exact p*<*0.05 with Bonferroni correction) enriched near invertons. We repeated this analysis independently for each inverton group. We also repeated this analysis by subsetting genes within the +/-5kb window to cases of (a) non-inverton-intersecting genes that were oriented with their 5’ end proximal to the inverton and were either adjacent to the inverton or all intervening genes were also oriented with their 5’ end proximal to the inverton such that a promoter in the inverton could potentially drive expression of said genes – i.e., regulatable, (b) non-inverton-intersecting genes that did not meet the criteria in (a), i.e., non-regulatable, and (c) genes that intersected a detected inverton.

### Invertase detection, categorization, and detection of links to inverton groups

Using PGAP annotations, we focused on extracting all coding sequences whose annotations contained mention of ‘invertase’, ‘integrase’, and ‘recombinase’. Treating these collectively as potential invertase genes, we used Clustal-omega [38] to perform a multiple sequence alignment on the translated amino acid sequences. We used TreeCluster to cluster invertases into distinct groups based on protein homology, using a tree-distance cutoff of 0.8 (Supplementary Section S6) to yield 176 invertase groups. For each invertase group, we counted for each of the 126 inverton groups the number of instances (i) an invertase in the invertase group is located within +/-5kb of an inverton in the inverton group, (ii) an invertase in another invertase group is located within +/-5kb of an inverton in the inverton group, (iii) an invertase in the invertase group is located within +/-5kb of an inverton in another inverton group, and (iv) an invertase in another invertase group is located within +/-5kb of an inverton in another inverton group. For each invertase-group / inverton-group pair, we compiled these four counts into a 2x2 contingency table, and identified significantly (Fisher’s Exact p*<*0.05 with Bonferroni correction) linked invertase-group / inverton-group pairs.

### Differential analysis of inverton orientation between different microbial growth conditions

To identify invertons whose orientation significantly differed between different microbial growth conditions A-vs.-B, we compared counts of forward and reverse supporting reads (Pe_F, Pe_R) pooled across all samples from condition A versus condition B. For each inverton we compiled the four resulting counts (forward reads pooled across condition A, reverse reads pooled across condition A, forward reads pooled across condition B, and reverse reads pooled across condition B) into a 2x2 contingency table, and applied a Fisherexact test to identify invertons whose forward-vs.-reverse counts significantly (p*<*0.05) linked to microbial growth condition, with Bonferroni correction to account for multiple hypothesis testing. We focused on two such A-vs.-B comparisons: (i) samples from pure strain isolate cultures vs. mouse stool samples to explore effect of community colonization in vivo, and (ii) samples from in vitro mucin-carrier vs. supernatant cultures.

### Differential gene expression predictions and validation in *Bacteroides thetaiotaomicron*

We predicted genes to be differentially expressed if they were located downstream of an inverton with a putative promoter motif, and the orientation of the associated inverton was also identified as linked to microbial growth conditions. We validated the 12 total such genes that were predicted to be differentially expressed between mouse stool and pure isolate culture in the *Bacteroides thetaiotaomicron* genome by analyzing published RNA-seq dataset from *Bacteroides thetaiotaomicron* that included both pure isolate culture as well as mouse stool samples [36]. We first used hocort [43] to remove mouse transcriptome derived reads from RNA-seq read libraries, then aligned to hCom2 coding sequences using bowtie2 [44]. Read counts were normalized using conditional quantile normalization with the cqn R package [45] to obtain estimates of reads per kilobase per million mapped reads (RPKM). We then calculate ratios of median RPKM between RNA-seq samples from pure isolate culture versus mouse stool samples to validate predicted differential expression.

#### Longitudinal analysis of inverton orientation across timepoints

To identify invertons whose orientation significantly differed between early vs. late generation mouse samples, we compared counts of forward and reverse supporting reads (Pe_F, Pe_R) pooled across all samples from mouse generation 1 [SC1] versus mouse generations 2-5 [SC2,SC3,SC4,SC5]. For each inverton we compiled the four resulting counts (forward reads pooled across generation 1 mice, reverse reads pooled across SC1 mice, forward reads pooled across [SC2,SC3,SC4,SC5] mice, and reverse reads pooled across [SC2,SC3,SC4,SC5] mice) into a 2x2 contingency table, and applied a Fisherexact test to identify invertons whose forward-vs.-reverse counts significantly (p*<*0.05) linked to microbial growth condition, with Bonferroni correction to account for multiple hypothesis testing. We then repeated this analysis with all four possible cutoffs for early vs. late generation ([SC1,SC2]-vs.-[SC3,SC4,SC5], [SC1,SC2,SC3]-vs.-[SC4,SC5], and [SC1,SC2,SC3,SC4]-vs.-[SC5]). Invertons were identified as having significant association with mouse generation timepoint if any of these tests passed the p*<*0.05 (with Bonferroni correction) threshold. This same approach was used to identify invertons whose orientation significantly differed between early vs. late passage *in vitro* mixed culture samples (testing ([P1]-vs.-[P2,P3,P4,P5,P6], [P1,P2]-vs.-[P3,P4,P5,P6], [P1,P2,P3]-vs.-[P4,P5,P6], [P1,P2,P3,P4]-vs.-[P5.P6], and [P1,P2,P3,P4,P5]-vs.-[P6]), independently analyzing mucin-carrier-attached and supernatant sample types.

### Data visualization

Custom python and R scripts were developed for data visualization, using the following packages: scipy [46], pandas [47], numpy [48], seaborn [49], python [50], jupyter note-book [51], statsmodels [52], biopython [53], matplotlib [54], R [55], ggtree [56], treeio [57], ggnewscale [58], phytools [59], tidyr [60], dplyr [61], stringr [62], ggplot2 [63], ape [64].

## Discussion

Here we presented a community-wide analysis of invertons in the defined gut microbial community hCom2, providing a customized workflow for detecting invertons in defined microbial communities based on the previously published PhaseFinder algorithm which we used to generate a comprehensive catalog of inverton locations across hCom2. Using the sequence homology found in the IR regions of these detected invertons, we categorized discovered invertons into groups, and used these groups to identify enriched motifs. This uncovered a number of promoter-like motifs – including some which were previously undescribed – whose instances were consistently found upstream of their nearest gene. Analyzing the proximity of invertons to nearby genes, we also revealed links between specific groups of invertons and specific groups of invertases, suggesting a potential regulatory link.

By analyzing large scale metagenomic sequencing of a defined community across multiple sample types, including isolate cultures vs. mouse fecal samples as well as surface-attached vs. liquid phase cultures, we were able to observe differences in inverton orientation probabilities associated with gut colonization and surface adhesion. By detecting directionally biased invertons in more than a third (53/125) of all strains in hCom2, we directly confirmed the hypothesis that colonization and adhesion are associated with inverton dynamics. A key advantage of performing this analysis in hCom2 was the availability of high quality genomes and genome annotations, which enabled us to leverage the discovery of directionally biased invertons into bioinformatic predictions of not only the identity of specific modulated genes, but also the direction of predicted modulation – for instance we predicted that colonization / adhesion are associated with a remodeling of EPS moieties in *Bacteroides*, aligning with previous work [9]. Beyond different sample types, we also identified invertons whose orientations exhibited changes over timepoints both *in vivo* (across mouse generations) and *invitro* (across culture passages). Our findings suggest active inverton dynamics across a range of timescales during surface adhesion and gut colonization, mediated by additional layers of regulatory control beyond IR sequence recognition alone.

We conclude by noting several limitations to our work and point to areas for further exploration. As our analyses are computational in nature, our predicted inverton-associated gene modulations, as well as links between invertase and invertons are only statistical associations. Mechanistic validation of these predictions will require future experimental work, for instance synthetic manipulations such as phase-locked inverton constructs to measure bacterial phenotypes when a particular inverton is locked in a forward or reverse orientation. These synthetic approaches can be combined with more sophisticated readouts such as metatranscriptomics on mixed bacterial communities to validate whether putative promoter motifs identified here indeed drive gene expression as we predict, expanding on the limited example we tested with *Bacteroides thetaiotaomicron* VPI-5482. Beyond promoters, future work should follow-up on additional motifs identified as enriched downstream of CDSs, to investigate whether they play any regulatory roles, for instance as transcriptional terminators. Additional experimental manipulations can also be used to target knockout or over-expression of predicted invertases, to test whether they indeed regulate inverton flipping of their predicted targets. Furthermore, the current analysis has focused on bacterial growth conditions with minimal environmental stress. Future work could augment the community by incorporating pathogenic taxa, and investigate how the presence of environmental stressors such as antibiotic exposure affect the inverton landscape. By obtaining more sequencing data in variable growth conditions, we would likely also expand the catalog of known invertons, given the observation that invertons flip in certain growth conditions. For instance, while we were only able to call 557 invertons using isolate culture sequencing reads during workflow benchmarking, by augmenting our analysis with sequencing data from different growth conditions (e.g., mixed in vitro culture and mouse stool samples), this increased to 1837 called invertons. Finally, while our work here has focused on inverton-mediated genetic variability, it would be valuable in the future to explore how inverton dynamics across microbial communities are linked to other forms of genetic variability, such as changes that arise as a consequence of mutation and horizontal gene transfer, to more comprehensively profile the cumulative effects of genetic variability on community function.

## Supplementary information

Supplementary Sections S1-S18 are available in the Supplementary Information PDF file. Supplementary Tables S1-S18 are included as CSV files and contain the following data:

Table S1: Metadata on hCom2 strains analyzed in this work

Table S2: Read libraries analyzed in this work with associated metadata

Table S3: Forward/reverse read counts of invertons in isolate culture comparing original PhaseFinder vs PhaseFinderDC

Table S4: Forward/reverse read counts of all invertons across all samples, determined using PhaseFinderDC

Table S5: Metadata on all identified invertons

Table S6: Enriched motifs detected across inverton IR sequences Table S7: Enriched motifs detected across full inverton sequences

Table S8: Instances of detected motifs from Table S7 across hCom2 genomes Table S9: Metadata on inverton-proximal genes across hCom2

Table S10: Enrichment of proximal genes by gene type, inverton group, regulatable-vs.-nonregulatable-vs.-intersecting

Table S11: Invertase genes across hCom2

Table S12: Enrichment of invertase groups proximal to inverton groups

Table S13: Directionally biased invertons comparing between (1) isolate culture / mouse stool and (2) carrier-attached / liquid-phase mixed culture samples

Table S14: Gene regulation predictions based on directionally biased invertons with promoter motifs

Table S15: Gene families enriched for inverton-based regulation

Table S16: Metadata on transcriptomic datasets used for gene regulation prediction validation

Table S17: Gene expression estimated based on transcriptomic datasets

Table S18: Directionally biased invertons comparing across timepoints (mouse generation and mixed culture passage)

## Acknowledgements.

We thank A. Lind, B. Smith, V. Dubinkina, C. Zhao, J. Wirbel, and M. Fischbach for helpful discussions on the manuscript. This work is supported by funding from Chan Zuckerberg Biohub, Burroughs Wellcome Fund, Gladstone Institutes, NSF grant #1563159, and NHLBI grant #HL160862.

Declarations

## Availability of data and materials

The mouse stool sequencing data generated in this study have been deposited in the NCBI database under BioProject accession code PRJNA1119053. In vitro cultures were analyzed using previously published data [27, 28], as well as new sequencing data that have been deposited in the NCBI database under BioProject accession code PRJNA1119029. Metadata for all metagenomic read libraries analyzed can be found in Supplementary Table S2. Key processed data generated in this study are provided in the Supplementary Tables.

PhaseFinderDC code available at: https://github.com/xiaofanjin/PhaseFinderDC

Code and additional data used for analysis and visualization available at: https://github.com/xiaofanjin/hcom2-invertons

## Author’s contributions

X.J., A.C., and K.S.P. contributed to the design and implementation of the research, to the analysis of the results and to the writing of the manuscript. F.B.Y., A.D., M.J., A.M.W., and J.Y. contributed to the implementation of the research. R.C. and A.S.B. contributed to the writing of the manuscript. All authors reviewed the manuscript.

## Notes

### Competing Interest Statement

The authors have declared no competing interest.

