## Supplementary Information for "Comprehensive profiling of genomic invertons in defined gut microbial community reveals associations with intestinal colonization and surface adhesion"

### S1 Characteristics of called invertons

Using Clustal-omega multiple sequence alignment [1], we generated a tree of all 1837 called invertons based on homology of their IR sequences, and used TreeCluster to categorize each of these invertons into one of 146 inverton groups – we further labelled each inverton by phylum and whether it intersects a gene or not (Fig. S1A). Many (especially larger), but not all inverton groups are dominated by invertons from a single bacterial phylum (Fig. S1B). The proportion of invertons that intersect a gene varies by phylum (Fig. S1C).

### S2 Optimizing tree clustering branch length T using MEME

To find the optimal tree clustering branch length cutoff T for generating inverton groups based on the MSA of their IR sequences, we tested across the full range of possible cutoffs (T=0 to T=1) in increments of 0.01 (i.e., testing T=0.01, 0.02, ... 0.99). For each tested value of T, we took the generated groups of invertons and used MEME to discover enriched motifs in each group of IR sequences. We then count the total number IR sequences for which a motif instance was discovered, and chose the optimal T to maximize this count (Fig. S2), yielding an optimal value of T=0.60.

### S3 Characteristics of top 29 inverton groups

We detected a total of 29 inverton groups for which at least 10 invertons were present from a single bacterial phylum. The top 6 of these inverton groups were dominated by invertons from Bacteroidota (groups 131, 126, 7, 136, 143) and *Akkermansia muciniphila* (group 138), and had IR sequence motifs (detected by running MEME on IR sequences from each group) that closely matched those of previously described Bacteroidota and *Akkermansia muciniphila* IR motifs from Jiang *et al.* [2] (Fig. S3). Four of the groups also contained promoter sequence motifs (detected by running MEME on full inverton sequences from each group) that closely matched previously described Bacteroidota and *Akkermansia muciniphila* promoter motifs [2] (Fig. S3)

Most of these top 29 inverton groups are dominated by a single bacterial phylum. For instance, inverton groups 128, 137, 122, 48, and 68 contain 23/28, 16/17, 12/16, 10/17, and 10/12 from Bacteroidota. We also found inverton groups dominated by phylum Firmicutes_A. For instance inverton group 139 contains 32/33 invertons from phylum Firmicutes_A spread across 17 distinct strains, while inverton group 125 contains 17/17 invertons from phylum Firmicutes_A, 15 of which derived from a single strain of *Blautia wexlerae* DSM-19850. An example of an exception is group 145 which contains representatives from both Bacteroidota and Firmicutes_A (Fig. S3, Supplementary Section S4).

### S4 Group 145 inverton characteristics

Inverton group 145 contains 20 invertons, 9 from phylum Bacteroidota (across 5 strains) and 10 from phylum Firmicutes_A (across 7 distinct strains). Remarkably, IR sequences for these invertons are highly conserved despite the evolutionary distance between their strains of origin – furthermore, IR sequences do not cluster by phylum (Fig. S4A).

Inverton group 145 invertons are furthermore characterized by an enrichment of intersections with coding sequences from the restriction endonuclease subunit S gene family (Fig. 3E,F), as well as and enrichment of nearby invertases from invertase group 96 (Fig. 4B) which are annotated by PGAP as tyrosine-type recombinase/integrase. Multiple sequence alignment of amino acid sequences of group 145 inverton intersecting restriction endonuclease subunit S gene family genes (Fig. S4B) as well as nearby tyrosine-type recombinase/integrase genes (Fig. S4C) revealed low and high degrees of sequence conservation respectively. Collectively, these findings suggest that group 145 invertons may potentially have spread across phyla via horizontal gene transfer mediated by tyrosine-type recombinase/integrase genes, conferring restriction enzyme genes with flexible specificity [3–6].

### S5 Discovery of promoter-like motifs

Using our promoter prediction approach (Methods – Motif detection and promoter prediction in invertons), we identified 8 putative promoter motifs whose instances across hCom2 were enriched in a consistent orientation upstream of the nearest gene (Fig. S5A-H). A motif found in the *Akkermansia muciniphila* dominated inverton group 138 – motif 138-2 – closely resembled the previously described *Akkermansia* motif [2] and also exhibited numerous promoter-like instances, but did not pass the $p<0.001$ significance cutoff (Fig. S5I). In addition to putative promoters, we also discovered 8 motifs whose instances were significantly enriched in a consistent orientation downstream their nearest gene (Fig. S5J-Q)

### S6 Grouping invertases based on sequence homology

We used multiple sequence alignment (with Clustal-omega) to build a tree of all invertase genes in hCom2, with genes being classified as invertase if their PGAP annotation included one of the following strings: 'invertase', 'recombinase', 'integrase'. Using TreeCluster [7] with cutoff T=0.80, we generated 176 groups of invertases based this tree of amino acid sequence similarity (Fig. S6)

### S7 Inverton group 131 is not enriched for nearby invertases

Out the three inverton groups for which we detected a motif similar to the previously described Bacteroidota promoter motif [2], groups 126 and 136 are both enriched for nearby invertases (from invertase groups 103 and 102 respectively, Fig. 4B,D,E). However, inverton group 131 is not enriched for nearby invertases (Fig. S6)

### S8 Invertons with dynamic F-vs.-R orientation across timepoints

A number of example invertons with shifting forward-vs.-reverse orientation trends over timepoints (either *in vitro* passages or *in vivo* mouse generations) were identified, with inversion ratios (number of reverse reads over total reads) plotted over time (Fig. 6). Here we additionally plot the same data as forward-vs.-reverse read count scatterplots, with accompanying genome diagrams of the regions surrounding each inverton for both *in vivo* (Fig. S8A) and *in vitro* (Fig. S8B) examples.

### S9 Group 138 *Akkermansia muciniphila* ATCC-BAA-835 inverton dynamics across mouse generations

Most of the invertons (32/39) from group 138 originated in *Akkermansia muciniphila* ATCC-BAA-835 – these invertons had highly conserved IR sequences (Fig. S9A), but varying dynamics across mouse generations with some but not all exhibiting time-dependent behavior (Fig. S9B,C) similar to the example highlighted in Fig. 6A.

### S10 Collection of invertons in *Clostridium sp.* D5 with similar *in vitro* dynamics

19 invertons *Clostridium sp.* D5 with low IR sequence conservation (Fig. S10A) all exhibited highly similar dynamics *in vitro*, characterized by low levels of reverse orientation on carrier-attached cultures with virtually no reverse orientation reads on corresponding supernatant cultuures (Fig. S10B,C).

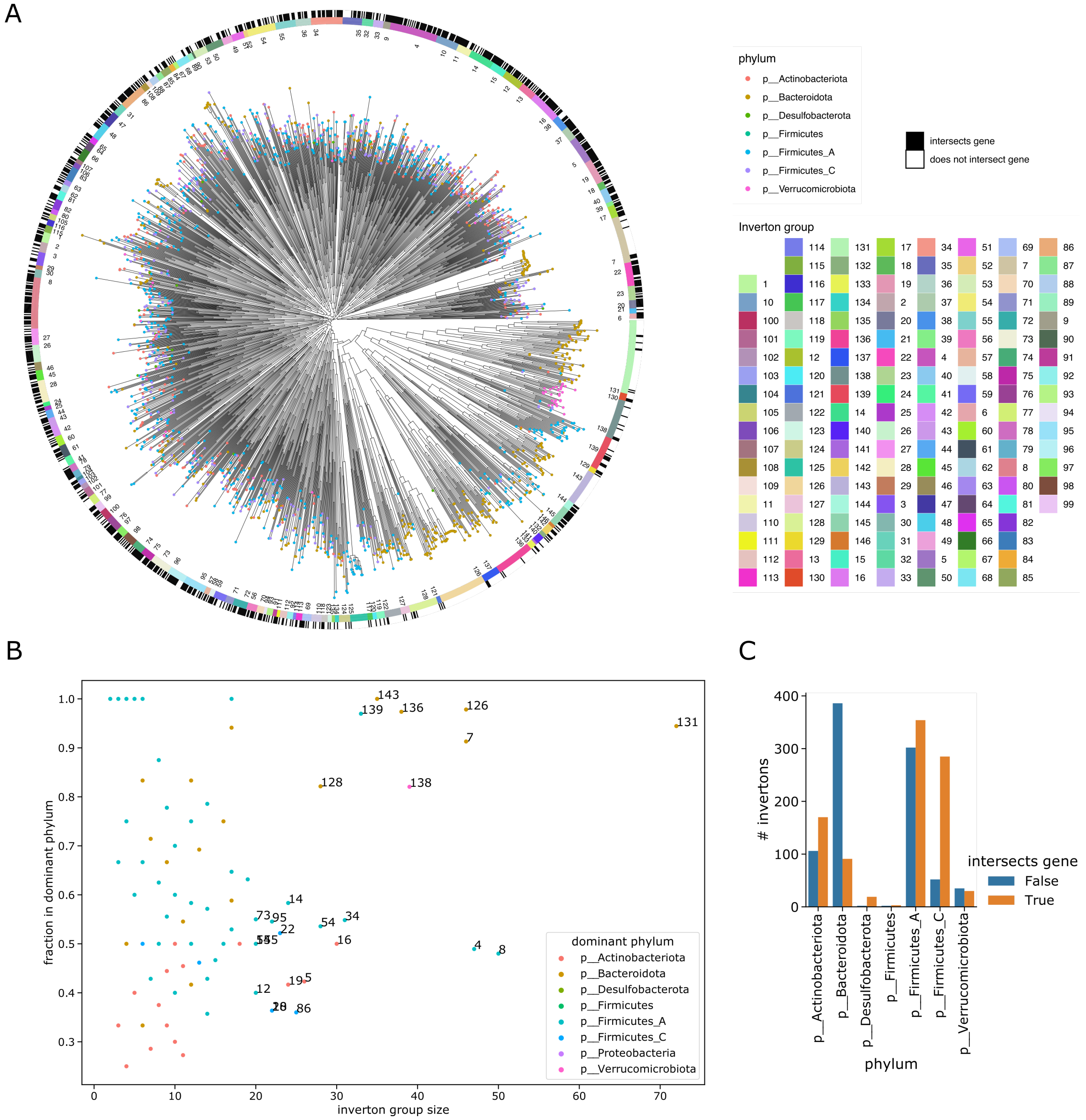


Figure S1: **Classifying invertons based on IR sequence homology, bacterial phylum, and gene intersection** **(A)** Tree of all 1837 invertons generated by MSA of IR sequences, labelled by bacterial phylum (tip colors), inverton group, (inner ring), and whether or not the inverton intersects a gene CDS (outer ring). **(B)** Scatterplot of inverton groups by number of invertons contained (horizontal axis) and fraction of invertons in dominant phylum (vertical axis), points colored according to dominant phylum in group, groups with 20 or more invertons are labeled. **(C)** Counts of invertons by bacterial phylum, colored by whether or not they intersect a gene CDS.

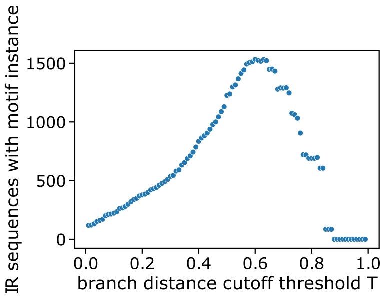


Figure S2: **Choosing optimal tree clustering branch length T to group invertons** Number of IRs with MEME discovered motif instance as a function of branch length T – highest count (1532/1837) occurs for T=0.60.

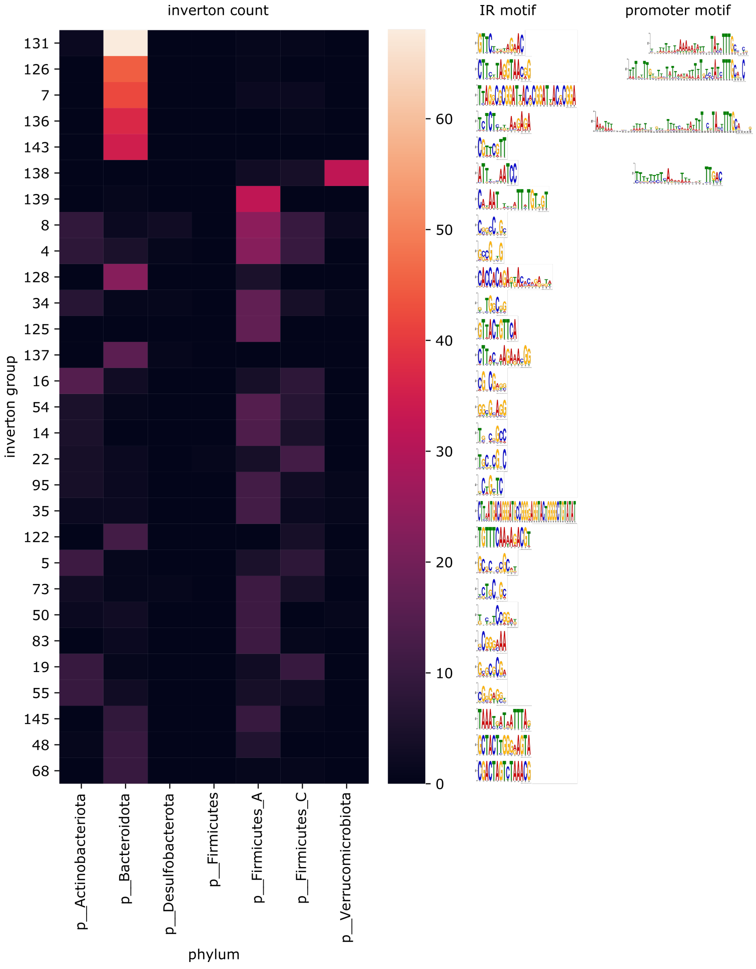


Figure S3: **Distribution of invertons in top 29 inverton groups** Inverton counts in top 29 inverton groups by phylum, each group labelled with respective IR motif. IR motifs from groups 131, 126, 7, 136, and 143 closely match Bacteroidota IR motifs 4, 3, 0, 1, and 2 from Jiang *et al.* respectively, while motif from group 138 closely matches their previously described *Akkermansia muciniphila* motif [2]. Groups 126, 131, 136 and 138 are further labeled with sequence motifs (motifs 126-2*, 131-2, 136-2* and 138-2 respectively, *denotes reverse complement) that closely match previously described Bacteroidota and *Akkermansia muciniphila* promoter motifs [2]

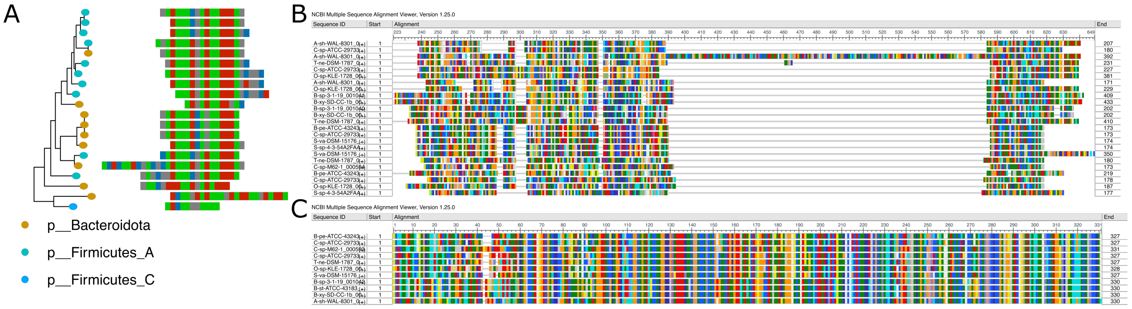


Figure S4: **Multiple sequence alignments associated with group 145 invertons (A)** Multiple sequence alignment of group 145 inverton IR sequences. **(B)** Multiple sequence (amino acid) alignment of restriction endonuclease subunit S gene family genes intersecting group 145 invertons. **(C)** Multiple sequence (amino acid) alignment of tyrosine-type recombinase/integrase genes near (<5kb) group 145 invertons.

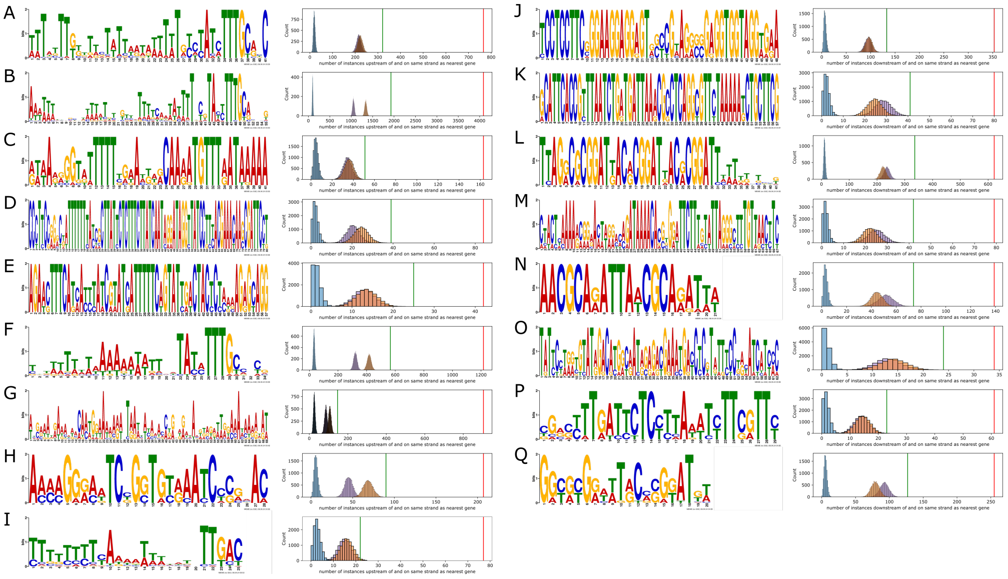


Figure S5: **Motifs enriched in consistent strand orientation rerlative to nearest gene (A)** Instances of motif 126-2 (reverse complement) were significantly enriched upstream of and in same strand as their nearest genes. This motif closely resembles a previously described Bacteroidota promoter motif [2]. Histogram colors as in Fig. 2E. **(B)** Instances of motif 136-2 (reverse complement) were significantly enriched upstream of and in same strand as their nearest genes. This motif closely resembles a previously described Bacteroidota promoter motif [2]. **(C)** Instances of motif 143-2 (reverse complement) were significantly enriched upstream of and in same strand as their nearest genes. **(D)** Instances of motif 68-2 (reverse complement) were significantly enriched upstream of and in same strand as their nearest genes. **(E)** Instances of motif 128-8 were significantly enriched upstream of and in same strand as their nearest genes. **(F)** Instances of motif 131-2 were significantly enriched upstream of and in same strand as their nearest genes. This motif closely resembles a previously described Bacteroidota promoter motif [2]. **(G)** Instances of motif 131-5 were significantly enriched upstream of and in same strand as their nearest genes. **(H)** Instances of motif 62-1 were significantly enriched upstream of and in same strand as their nearest genes. **(I)** Instances of motif 138-2 were enriched – but not significantly ($p < 0.001$) so – upstream of and in same strand as their nearest genes. This motif closely resembles known *Akkermansia* promoter motif [2]. **(J)** Instances of motif 3-1 (reverse complement) were significantly enriched downstream of and in same strand as their nearest genes. **(K)** Instances of motif 51-2 (reverse complement) were significantly enriched downstream of and in same strand as their nearest genes. **(L)** Instances of motif 7-1 (reverse complement) were significantly enriched downstream of and in same strand as their nearest genes. **(M)** Instances of motif 126-5 were significantly enriched downstream of and in same strand as their nearest genes. **(N)** Instances of motif 127-1 were significantly enriched downstream of and in same strand as their nearest genes. **(O)** Instances of motif 48-2 were significantly enriched downstream of and in same strand as their nearest genes. **(P)** Instances of motif 48-4 were significantly enriched downstream of and in same strand as their nearest genes. **(Q)** Instances of motif 7-2 were significantly enriched downstream of and in same strand as their nearest genes.

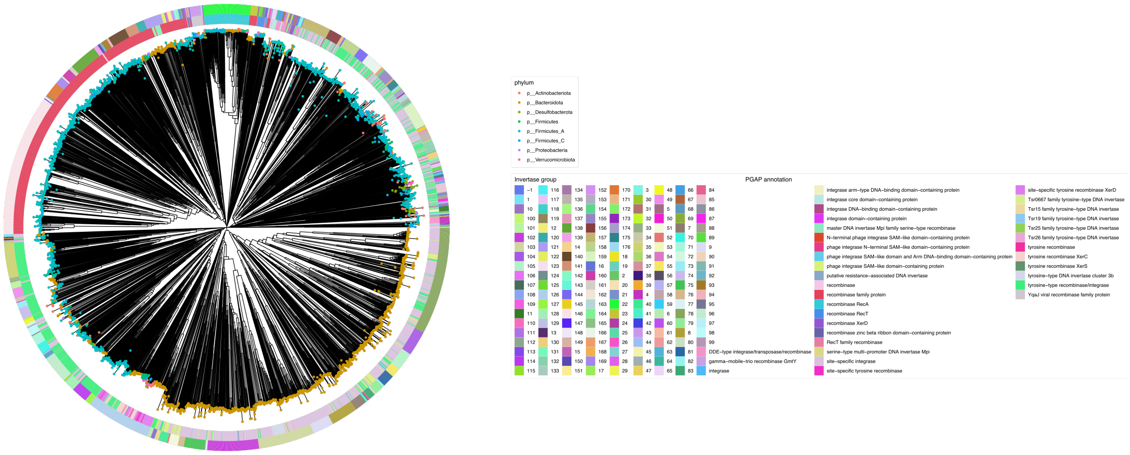


Figure S6: **Tree of invertase amino acid sequence similarity** Tree of hCom2 invertase gene sequences, tip labels represent bacterial phylum, outer ring colors represent invertase group, inner ring colors represent PGAP annotation.

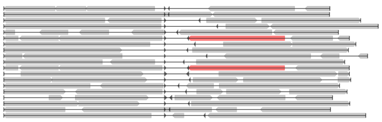

Figure S7: **Lack of invertases near inverton group 131 invertons** Genome diagram of regions surrounding group 131 invertons, with invertase group 102 genes highlighted in red – for simplification if multiple invertons of this group are found in the same genome, a single representative is plotted. Note markedly fewer invertases surrounding group 131 invertons compared to groups 126, 136 (Fig. 4D,E).

A

B

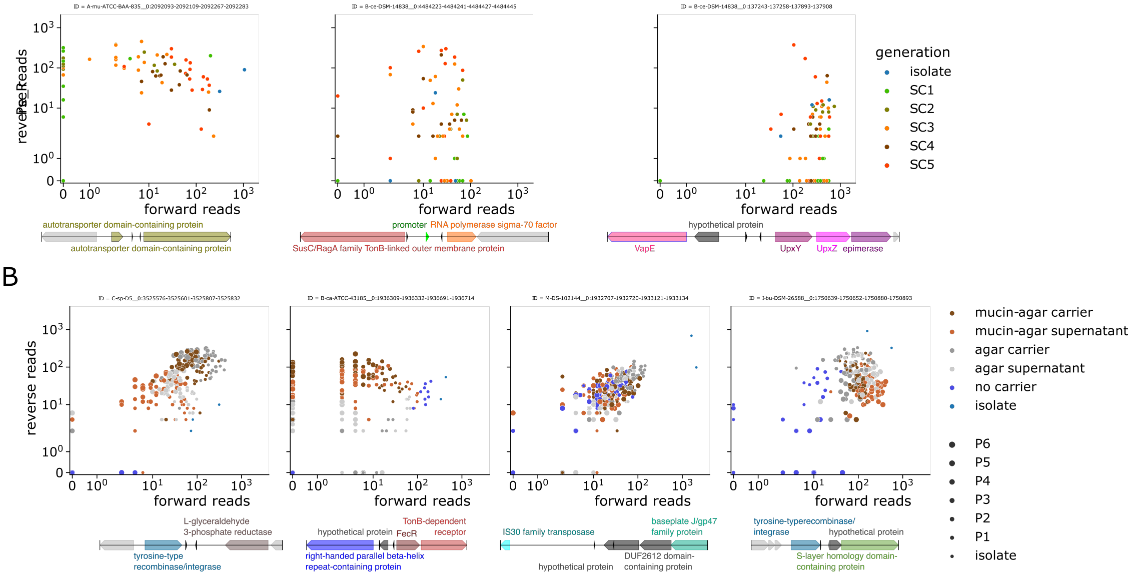


Figure S8: **F-vs.-R scatterplots and genome diagrams of invertons with time-dependent behavior (A)** Forward vs. reverse read count scatterplots of invertons highlighted in Fig. 6A-C, with accompanying genome diagrams of surrounding regions. **(B)** Forward vs. reverse read count scatterplots of invertons highlighted in Fig. 6D-G, with accompanying genome diagrams of surrounding regions. Note these plots include three additional *in vitro* culture conditions (plain agar carriers, plain agar supernatant, no carrier controls [8]) that were omitted for simplicity in Fig. 6D-G.

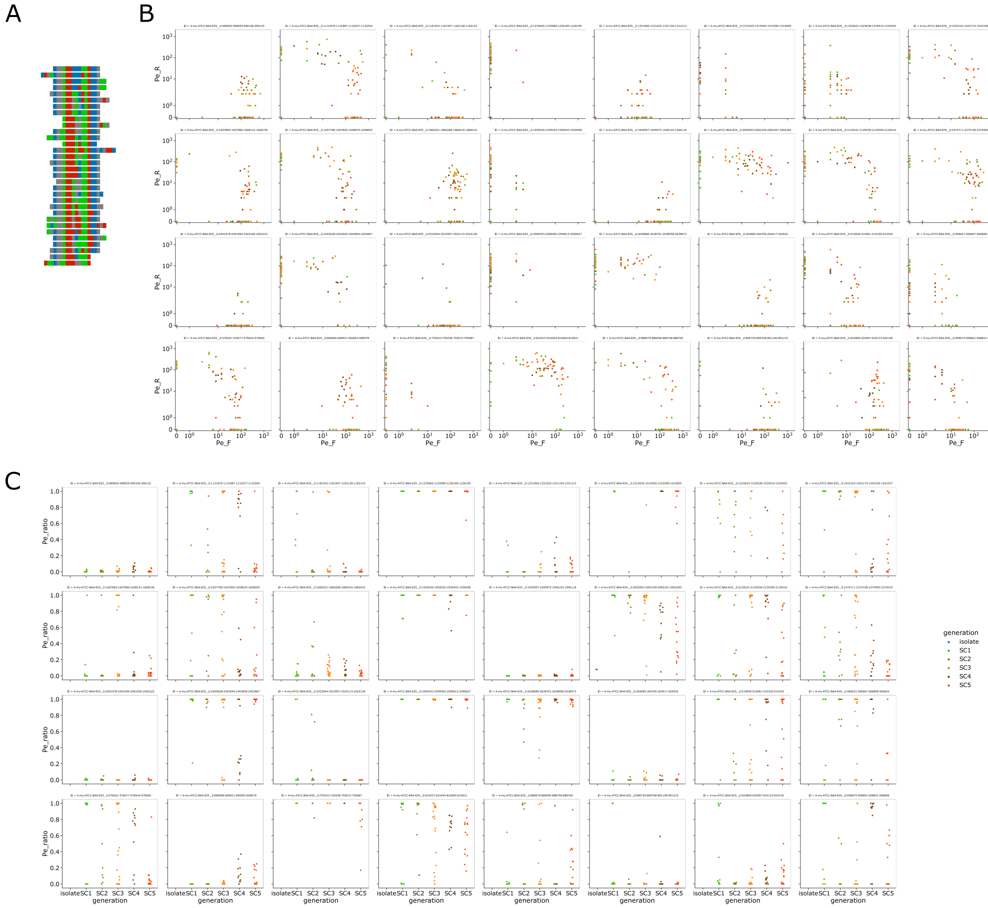

Figure S9: **Dynamics of group 138 *Akkermansia muciniphila* ATCC-BAA-835 invertons (A)** Multiple sequence alignment of IR sequences of group 138 invertons from *Akkermansia muciniphila* ATCC-BAA-835. **(B)** Forward vs. reverse read count scatterplots of group 138 invertons from *Akkermansia muciniphila* ATCC-BAA-835, colored by mouse generation. **(C)** Inversion ratio (R/(R+F) across mouse generations plotted for group 138 invertons from *Akkermansia muciniphila* ATCC-BAA-835.

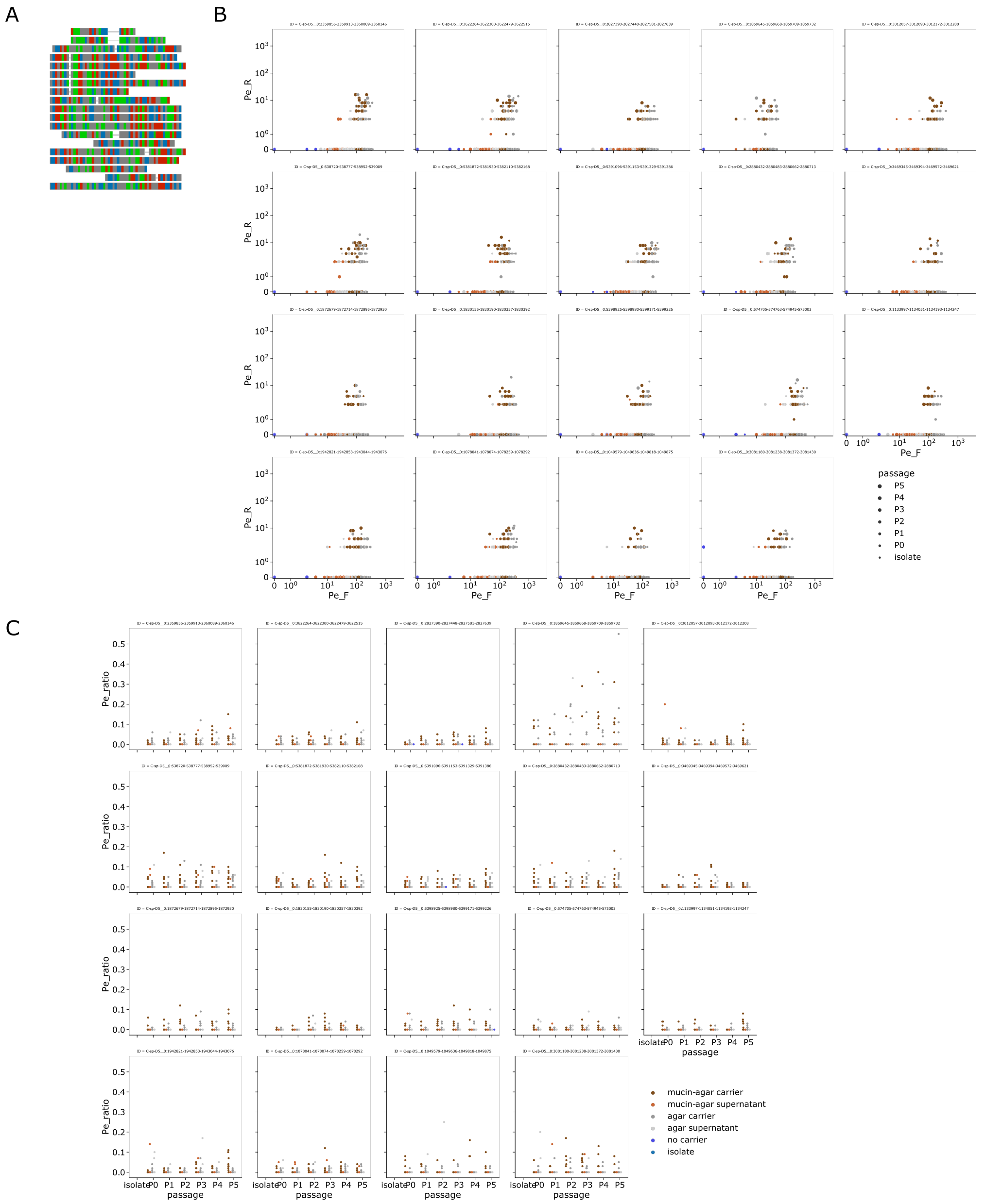

Figure S10: **Collection of *Clostridium sp.* D5 invertons with similar *in vitro* dynamics (A)** Multiple sequence alignment of IR sequences of 19 *Clostridium sp.* D5 invertons with similar *in vitro* dynamics. **(B)** Forward vs. reverse read count scatterplots of 19 *Clostridium sp.* D5 invertons with similar *in vitro* dynamics, colored by *in vitro* culture type. Note these plots include three additional *in vitro* culture conditions (plain agar carriers, plain agar supernatant, no carrier controls [8]) that were omitted for simplicity in Fig. 6D-G. **(C)** Inversion ratio (R/(R+F) across culture passage timepoints plotted for 19 *Clostridium sp.* D5 invertons with similar *in vitro* dynamics, colored by *in vitro* culture type. Note these plots include three additional *in vitro* culture conditions (plain agar carriers, plain agar supernatant, no carrier controls [8]) that were omitted for simplicity in Fig. 6D-G.
